# MuGVRE. A virtual research environment for 3D/4D genomics

**DOI:** 10.1101/602474

**Authors:** Laia Codó, Genís Bayarri, Javier Alvarez Cid-Fuentes, Javier Conejero, Adam Hospital, Romina Royo, Dmitry Repchevsky, Marco Pasi, Athina Meletiou, Mark D. McDowall, Fatima Reham, José A. Alcantara, Brian Jimenez-Garcia, Jürgen Walther, Ricard Illa, François Serra, Michael Goodstadt, David Castillo, Satish Sati, Diana Buitrago, Isabelle Brun-Heath, Juan Fernandez-Recio, Giacomo Cavalli, Marc Marti-Renom, Andrew Yates, Charles A. Laughton, Rosa M. Badia, Modesto Orozco, Josep Ll. Gelpí

**Affiliations:** Barcelona Supercomputing Center, Barcelonar, Spain; Institute for Research in Biomedicine, the Barcelona Institute of Science and Technology. Barcelona, Spain; Sch. of Pharmacy and Centre for Biomolecular Sciences, Nottingham, UK; European Molecular Biological Laboratory, European Bioinformatics Institute, Wellcome Genome Campus, Hinxton, UK; CNAG-CRG, Centre for Genomic Regulation (CRG), Barcelona Institute of Science and Technology (BIST), Baldiri i Reixac 4, 08028 Barcelona, Spain. Gene Regulation, Stem Cells and Cancer Program, Centre for Genomic Regulation (CRG), Dr. Aiguader 88, 08003 Barcelona, Spain. Universitat Pompeu Fabra (UPF), Barcelona, Spain; Institute of Human Genetics, CNRS, Univ. Montpellier, Montpellier, France; ICREA, Pg. Lluís Companys 23, 08010 Barcelona, Spain; Dept. of Biochemistry and Molecular Biomedicine, University of Barcelona, Barcelona, Spain

## Abstract

Multiscale Genomics (MuG) Virtual Research Environment (MuGVRE) is a cloud-based computational infrastructure created to support the deployment of software tools addressing the various levels of analysis in 3D/4D genomics. Integrated tools tackle needs ranging from high computationally demanding applications (e.g. molecular dynamics simulations) to high-throughput data analysis applications (like the processing of next generation sequencing). The MuG Infrastructure is based on openNebula cloud systems implemented at the Institute for research in Biomedicine, and the Barcelona Supercomputing Center, and has specific interfaces for users and developers. Interoperability of the tools included in MuGVRE is maintained through a rich set of metadata allowing the system to associate tools and data in a transparent manner. Execution scheduling is based in a traditional queueing system to handle demand peaks in applications of fixed needs, and an elastic and multi-scale programming model (pyCOMPSs, controlled by the PMES scheduler), for complex workflows requiring distributed or multi-scale executions schemes. MuGVRE is available at https://vre.multiscalegenomics.eu and documentation and general information at https://www.multiscalegenomics.eu. The infrastructure is open and freely accessible.

## INTRODUCTION

While major milestones have been achieved in determining the sequence of DNA, understanding its 3D folding, the connection between chromatin structure and genome functionality and the links between changes in chromatin structure and pathology are still major challenges that are attracting a large research effort and have created a new area of knowledge: 3D/4D genomics. Opposite to traditional sequencing projects, 3D/4D genomics faces a major problem related to the diversity of data types and formats generated, the variety of analysis methods, and the multi-resolution nature of the navigation in a multi-scale data space. Data used in the field range from simple sequence (1D genomics), typically enriched by functional and structural annotations (2D Genomics) to contact maps, single or multiple structures (at a very wide range of resolutions) and images.

ENA (1), EGA (2) and Ensembl (3) are the most common sources of 1D genomic data in Europe, whose contents are visualized thanks to tools such as “Genome browsers” (4-8) that provide an integrated view of both sequence and annotations (2D genomics). Many research infrastructures deal with this level of data, Galaxy (9), being by far the most popular.

A second level of data includes nucleic acid and protein structures determined at the atomistic level by X-Ray crystallography or NMR spectroscopy and deposited in the Protein Data Bank (10). Analysis and visualization of data at this level have, in practice, little overlap with the 2D genomics level. This represents a major caveat for people interested in specific DNA-protein complexes. An additional type of data includes simulation trajectories, i.e. a set of structures defining the conformational ensemble of DNA (and associated proteins), which is typically obtained through atomistic or coarse-grained molecular dynamics (MD) simulations (11-17). Specific databases on DNA simulation data (18) or protein-DNA complexes (19, 20), and tools to perform flexibility analysis of nucleic acids like NAFlex (21) or 3DNA (22) are available. Few tools allow mapping 3D structures onto the genomic sequences (23-25).

A third level of data is represented by nucleosome and protein positioning studies (26-30). Raw data is obtained from sequencing analysis (MNAse-seq, Chip-seq) and is usually limited to the annotation of specific binding sites. Data at this level is available from public repositories like ArrayExpress (31), where it is presented in a 1D manner without any connection to the 3D structure and flexibility of the chromatin fiber. At the upper end of the scale the main experimental strategies include FISH (32), and chromosome conformation capture (3C-like techniques (33)), which provide chimeric sequences representing interactions within the genome, from which structural insight can be derived after a complex manipulation of the data. Data visualization at this level is complex (34); several tools are already available (35-38), but they ignore any atomistic detail, or even the general description of the nucleosome string.

In summary, 3D (structure) and 4D (dynamics) genomics cover a large variety of data types and experimental and theoretical strategies. Tools and data repositories exist; but they are not integrated. As a result, research at the different levels is performed in isolation, and researchers almost ignore alternative views coming from different levels. The Multiscale Complex Genomics project (MuG, http://www.multiscalegenomics.eu/MuG) is one of the initiatives aiming to bridge the gap among the different levels of chromatin study and provide such an integrated view. Here, we present the MuG Virtual Research Environment (MuGVRE) infrastructure, designed to provide researchers with a single access point where data and tools covering the full spectrum, from sequence and atomistic data to chromosome capture results, can be used and combined to obtain a holistic picture of chromatin. MuGVRE is cloud based, allowing for an easy deployment and extension at the technical level. It is a base infrastructure where additional tools can be plugged-in to extend the functionality. The initial offer of tools already hosted, includes sequence, structure and Hi-C data analysis and visualizers; tools to handle nucleosome positioning, and tools for performing and analyzing simulation data from atomistic to coarse-grained levels. All the tools are accessible through an intuitive personal workbench, where most technical decisions are taken automatically by the system, allowing the user to concentrate on the scientific aspects of the analysis. We believe that MugVRE is the missing element to integrate the different views of physiological DNA.

## INFRASTRUCTURE DESIGN AND COMPONENTS

The MuGVRE infrastructure has been designed to fulfil the following principles:

### Technical

1. <u>Flexible structure</u>, able to adapt to the specific needs of the analysis tools, both in terms of software requirements, and computational resources.
2. <u>Software scheduler(s)</u>, able to manage analysis workflows, and computational resources in a transparent and adaptable manner. This should constitute an elastic infrastructure with automatic adaptation to user loads.
3. <u>Multi-scale execution</u>. Analysis workflows should be executed either at the cluster level, in HPC environments, or distributed infrastructures like EGI (https://www.egi.eu), and eventually in the forthcoming European Open Science Cloud (EOSC) ecosystem.

### Usage

4. <u>Web-based access</u> centered on the user workspace and complemented by full programmatic access using well-established interfaces.
5. <u>Data should be kept private</u>, through the appropriate Authentication and Authorization Infrastructure applied to all data transactions.

Supplementary Figure S1 shows a general schema of the computational infrastructure underlying MuGVRE.

## MuGVRE Main components

### Cloud deployments

MuGVRE infrastructure has been designed as a fully virtualized environment. At its present state, MuGVRE has been deployed in at the Starlife cloud infrastructure (https://www.bsc.es/marenostrum/star-life), at the Barcelona Supercomputing Center, using OpenNebula (https://opennebula.org/) and the KVM hypervisor (https://www.linux-kvm.org).

### Process management

MuGVRE uses two complementary layouts for process management: i) Sun Grid Engine (SGE, https://sourceforge.net/projects/gridscheduler/), in combination with OneFlow (https://docs.opennebula.org/5.4/advanced_components/application_flow_and_auto-scaling/appflow_use_cli.html), a component of the OpenNebula framework that allows managing Multi-VM applications and auto-scaling. SGE is used to manage applications where no complex workflows are necessary, requiring only to deploy additional workers on peaks of demand, ii) the COMPS Superscalar (COMPSs) programming model (and its python binding pyCOMPSs (39)), managed by the Programming Model Enactment Service (PMES) (40), which interacts with cloud infrastructures through Open Cloud Computing Interface (OCCI, http://occi-wg.org/) servers. PMES/pyCOMPSs are used to control complex workflows and distributed execution.

### Database manager

Operational data and metadata regarding installed tools, public repositories, and user’s files are maintained in a MongoDB database (https://www.mongodb.com). The MongoDB server also contains reference data as a copy of Protein Data Bank (10), Uniprot (41), and BiGNASim (18) database. User’s data is stored in a standard filesystem in its original format.

### Authentication and authorization system

Data privacy is maintained using the authentication and authorization server Keycloak (http://www.keycloak.org/) to handle all internal communications and user access. Keycloak implements OpenID Connect which allows for Web access on the code authorization flow of OAuth2, and a token-based authentication for the REST services. For registered users, authentication schemes based on username/password, but also third-party identity providers (Google, LinkedIn, Elixir) are accepted.

See Supplementary Material Section 1 for additional information about MuGVRE software components.

## USER WORKSPACE

### User access

MuGVRE can be accessed without authentication. Users are granted a private workspace to hold data and analysis results. Data is maintained for one week after the last access and can be recovered during this period through a unique URL address. Users desiring a longer interaction with MuGVRE are advised to register to get a permanent workspace. In such cases the user space is maintained up to two months after the last access.

### Personal workspace

The MuGVRE personal workspace is the central environment for user activity. It is based on a filesystem-based layout (see Figure 2), where both uploaded data and analysis results are available. Uploaded data should be annotated to specify data types and formats. This allows the MuGVRE workspace to offer an adapted toolkit for each file, including only compatible tools and visualizers (see Suppl. Material Section 2 for a description of the procedure, and Suppl.Table S3 for a description of accepted data types and formats). The user workspace has been laid out to provide an intuitive look-and-feel. The workspace itself is structured in projects, to keep data logically organized. Within each project the input data is distributed in folders: *Uploads* (uploaded data), and *Repository* (data obtained from public repositories). The remaining folders correspond to analysis operations (a new folder is generated for any new process started in the VRE). File lists can be filtered by any of the fields (name, format, data type, or project). Additionally, a tool-based filter is available to select only valid input files for the given tool.

**Figure 1.**
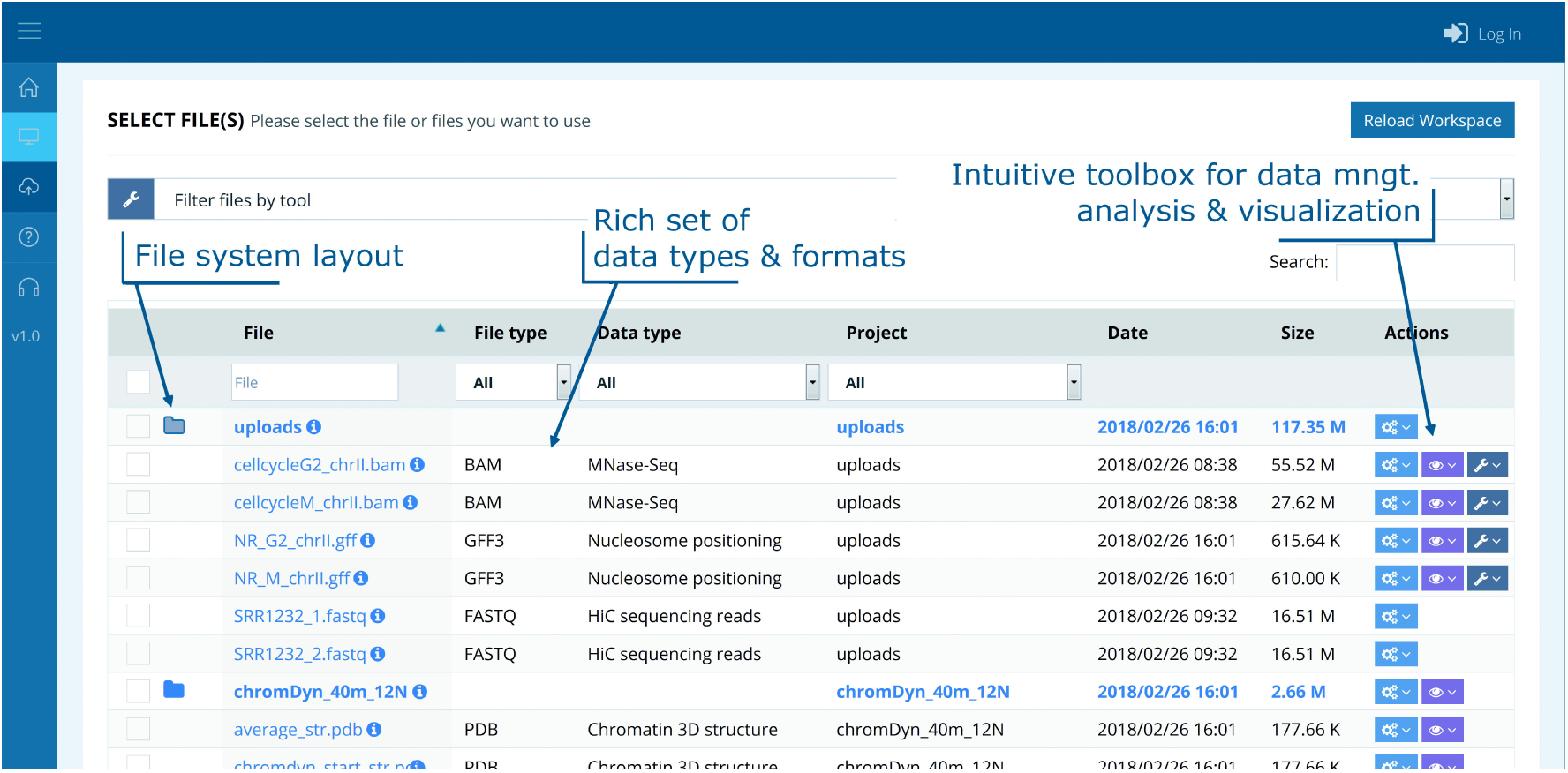
Screenshot of MuGVRE personal workspace

**Figure 2.**
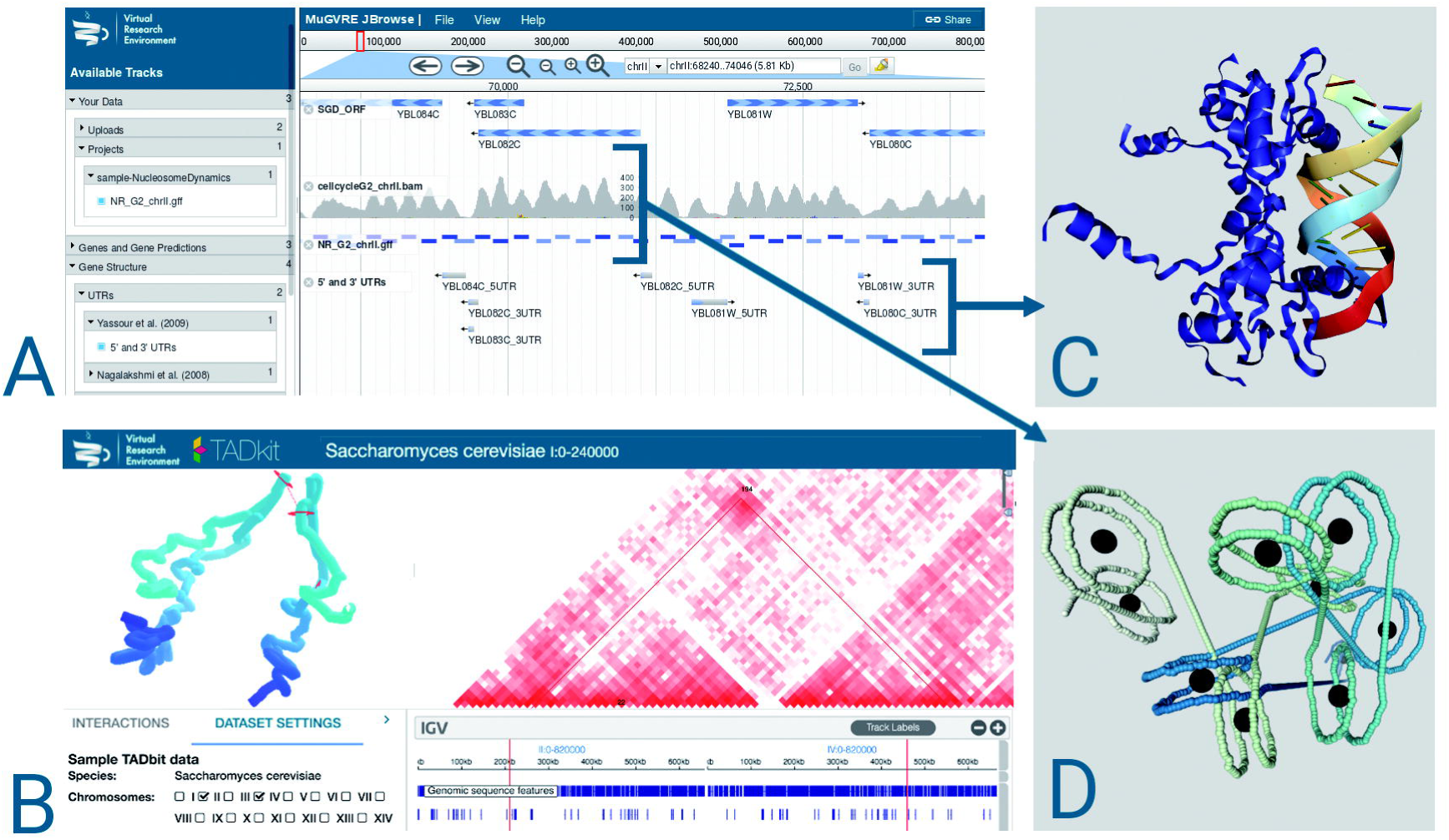
Sample visualizations at MuGVRE. A: Genome browser showing reference sequences, annotations, including a nucleosome positioning track (blue marks) (Jbrowse). B: Transcription Factor structure bound to cognate DNA fragment (NGL). C. HiC adjacent matrix, combined with genomics annotations, and a 3D model of chromatin structure (Tadkit), and D: Coarse-grained model of DNA fiber with several attached nucleosomes, from ChromatinDynamics (NGL).

Three interactive toolkits containing the following options are available:

- File toolkit: Download, rename, move, compress, delete files and folders, edit their metadata, rerun jobs.
- Visualization toolkit: Available visualizers for the specific file/s selection (based on their data type and format).
- Tools toolkit: Available tools for the specific file/s selection (based on their data type and format).

### Tools based access

In addition to the data centric approach, more experienced user may prefer to access directly to the analysis operation. To this end, user may select the desired operation from an ontology, covering all the operations provided by the installed tools. The user should then select the desired tool and fill in the required parameters. At this point, a selection of input files filtered out from the user workspace are suggested according to the metadata accompanying both, user’s data and tool input restrictions.

### Applications and developer access

MuGVRE is designed as an infrastructure open to any application designed for the analysis of 3D/4D genomics data. Tools installed in MuGVRE should accept free, unrestricted usage. Guidelines are available for developers (see Suppl. Material, Section 2). Developers wishing to include their applications are granted access to a specific workspace to manage tool definitions and execution details, and to edit the tool’s help pages, and perform execution tests. Also developers get access to usage statistics, execution logs, and job associated files and metadata. The current offer of tools is detailed in Suppl. Table S4. Several data visualizers are available (Suppl, Table S5). Figure 2 shows a representative set of the types of data than can be visualized in the infrastructure. JBrowse (5) is used for sequence data (see Figure 2A). In the figure, data from a Nucleosome Dynamics analysis of MNase-seq data is depicted (blue blocks for nucleosome calls, and coverage plot). The NGL visualizer (42) is used for 3D atomistic (see Figure 2B for a transcription factor-DNA complex) or coarse-grained structure data (output from Chromatin Dynamics, Figure 2D, depicting the structure of a fragment of chromatin fiber containing nucleosomes at positions shown in Fig. 2A). Finally, the TadKit visualizer (http://3DGenomes.org/tadkit, Figure 2C) allows the combination of 1D to 3D information taken from chromosome capture data.

## USAGE

Users can populate the MuGVRE workspace in several ways:

- *Direct upload into the workspace*, using the HTTPS protocol
- *Create files using an embedded text editor*. This is intended for data or metadata of reduced size.
- *Upload from an External URL*: MuGVRE can access external sites to download. This is the recommended procedure to obtain data from public repositories, or import bulky data, as the upload process becomes a batch job. HTTP and FTP protocols are accepted, also when user credentials are required.
- *From repository:* Data imported from the list of public repositories made browsable at the infrastructure (currently ArrayExpress (31) and BigNASim(18)).
- From sample data: selected input and output examples for the available tools are provided as help to start using the interface.

Files can be selected anywhere in the workspace, and added to the execution list, where a specific tool should be then selected. Alternatively, the user can select first an analysis tool from a list of available operations. Either selection mode opens a configuration screen where the user can assign data files (Suppl. Figure S7) to the appropriate input parameters, define additional settings, and launch the tool. Executions are performed in the background and do not require the user to keep the interactive session open. Results of the analysis are added to the workspace under a separate folder that contains the output files generated by the tool, log files, and a customized results page (Suppl. Figure S10). For a complete usage example, see Supplementary Material Section 3, where we show a series of screenshots of a session centered on the analysis of MNase-seq data from yeast Chr II on phases M and G2 of the cell cycle (data taken from (43)), analyzed with the Nucleosome Dynamics tool.

## DISCUSSION

3D/4D Genomics is an emerging field originated from the unplanned aggregation of different disciplines which have developed their tools, and associated data types and formats, independently. This diversity is a major obstacle towards the generation of a complete picture of chromatin structure and dynamics. MuGVRE has been designed as an integration space. It follows the traditional concept of the personal workbench, already used in general genomic workbenches like Galaxy (9) or GenePattern (44), or spaces designed for simulation data analysis like NAFlex (21). In this kind of environment, data and tools are available and the user has the freedom to design his/her own analysis pipelines. MuGVRE has an initial offer of tools and visualizers covering all levels of resolution, from atomistic simulation to chromatin fiber simulation, or Hi-C data analysis. Tools and visualizers are offered in a single space where, for example, chromosome conformation capture or nucleosome positioning data can be visualized along with sequence annotations and the structures and binding modes of the transcription factors affecting the same DNA region. A strong commitment of MuGVRE design is to free the user from understanding the technical side of the infrastructure. With this aim, not only the computational layout is hidden, but also most technical decisions are taken automatically by the system. For example, we have designed a comprehensive ontology of data types and formats that are checked internally to configure the options offered to the user. The user can just select a tool, and the workspace selects those files that match its input requirements. Output from the analyses are reusable following the same philosophy. As a result, the user can easily configure a pipeline taking only scientific decisions and not bothering about technicalities.

MuGVRE has been designed as a large and sustainable infrastructure, which relies now on the computational capabilities of the Barcelona Supercomputing Center (http://www.bsc.es), but technical decisions have been taken to assure the compatibility with other infrastructures like Elixir computational platforms (http://elixir-europe.org), EGI (https://www.egi.eu), or EUDAT (https://eudat.eu). The choice of a fully flexible cloud system, controlled with a multiscale software scheduler, and linked to HPC facilities, assures the usage of the optimal computational environment for each specific analysis.

MuGVRE is presented as an open platform with the aim of growing in functionality, since new tools can be easily incorporated by external developers. Hosting at MuGVRE can be an option for developers alternative to build a dedicated web site to run their tools.

MuGVRE was presented to the multiscale genomics community in November 2017, and has performed already over 6,500 analysis runs.

MuGVRE is a unique tool that aims to help researchers in the 3D/4D genomics field to gain an integrated view of discipline, sharing data among the diverse analysis levels and providing a complete and integrated view on DNA. We hope that MuGVRE will foster the development, deployment and use of new strategies for the analysis of the chromatin structure that were not envisioned simply because data was kept in separate silos.

## AVAILABILITY

MuGVRE: https://vre.multiscalegenomics.eu

General information and documentation: https://www.multiscalegenomics.eu

## ACKNOWLEDGEMENT

We are indebted to the entire MuG consortium and to the *β*-testers of the application for suggestions and comments.

