## Supplementary Material for "MuGVRE. A virtual research environment for 3D/4D genomics"

Laia Codó[1], Genís Bayarri[2], Javier Alvarez Cid-Fuentes[1], Javier Conejero[1], Adam Hospital[2], Romina Royo[1], Dmitry Repchevsky[1], Marco Pasi[3], Athina Meletiou[3], Mark D. McDowall[4], Fatima Reham[4], José A. Alcantara[2], Brian Jimenez-Garcia[1], Jurgen Walther[2], Ricard Illa[2], François Serra[5], Michael Goodstadt[5], David Castillo[5], Satish Sati[6], Diana Buitrago[2], Isabelle Brun-Heath[2], Juan Fernandez-Recio[1,7], Giacomo Cavalli[6], Marc Marti-Renom[5,8], Andrew Yates[4], Charles A. Laughton[3], Rosa M. Badia[1], Modesto Orozco[2,8], Josep Ll. Gelpí[1,8]*

1. Barcelona Supercomputing Center, Barcelona, Spain,
2. Institute for Research in Biomedicine, the Barcelona Institute of Science and Technology. Barcelona, Spain,
3. Sch. of Pharmacy and Centre for Biomolecular Sciences, Nottingham, UK,
4. European Molecular Biological Laboratory, European Bioinformatics Institute, Wellcome Genome Campus, Hinxton, UK
5. CNAG-CRG, Centre for Genomic Regulation (CRG), Barcelona Institute of Science and Technology (BIST), Baldiri i Reixac 4, 08028 Barcelona, Spain. Gene Regulation, Stem Cells and Cancer Program, Centre for Genomic Regulation (CRG), Dr. Aiguader 88, 08003 Barcelona, Spain. Universitat Pompeu Fabra (UPF), Barcelona, Spain.
6. Institute of Human Genetics, CNRS, Univ. Montpellier, Montpellier, France.
7. ICREA, Pg. Lluís Companys 23, 08010 Barcelona, Spain,
8. Dept. of Biochemistry and Molecular Biomedicine, University of Barcelona, Barcelona, Spain.

**CONTENTS**

1. **MuGVRE INFRASTRUCTURE AND COMPONENTS**
2. **PROCEDURE FOR THE MANAGEMENT OF TOOLS IN MUGVRE**
3. **USAGE EXAMPLE. NUCLEOSOME DYNAMICS ANALYSIS ON MNase-SEQ DATA FROM YEAST CHRII ON PHASES M AND G2 OF THE CELL CYCLE**
4. **REFERENCES**

### INDEX OF SUPPLEMENTARY TABLES

**Table S1.** Schema and example of Tool definition configuration file, required for registering a new tool

**Table S2.** Examples for the configuration files exchanged between VRE and tool VMs during the tool life cycle execution

**Table S3.** Data types and formats

**Table S4.** Analysis and Simulation tools

**Table S5.** Data visualizers available at MuG VRE

### INDEX OF SUPPLEMENTARY FIGURES

**Figure S1.** Scheme of MuGVRE computational infrastructure.

**Figure S2**. COMPSs master-worker architecture.

**Figure S3**. Life cycle of a tool execution in VRE, and how the information is transferred from the VRE user to the virtualized tools, and back to the VRE user.

**Figure S4**. Usage Example. Validation Form required to add the necessary metadata to uploaded files.

**Figure S5.** Usage Example. Workspace with toolkits

**Figure S6.** Usage Example. Jbrowse view of coverage of input BAM files

**Figure S7**. Usage Example. Parameter input and submit

**Figure S8**: Usage Example. Job progress display. Digested view of the application log

**Figure S9**. Usage Example. Project folder including output files

**Figure S10.** Usage Example. Summary results page

**Figure S11.** Usage Example. JBrowse view of Nucleosome analysis results

### INFRASTRUCTURE AND COMPONENTS

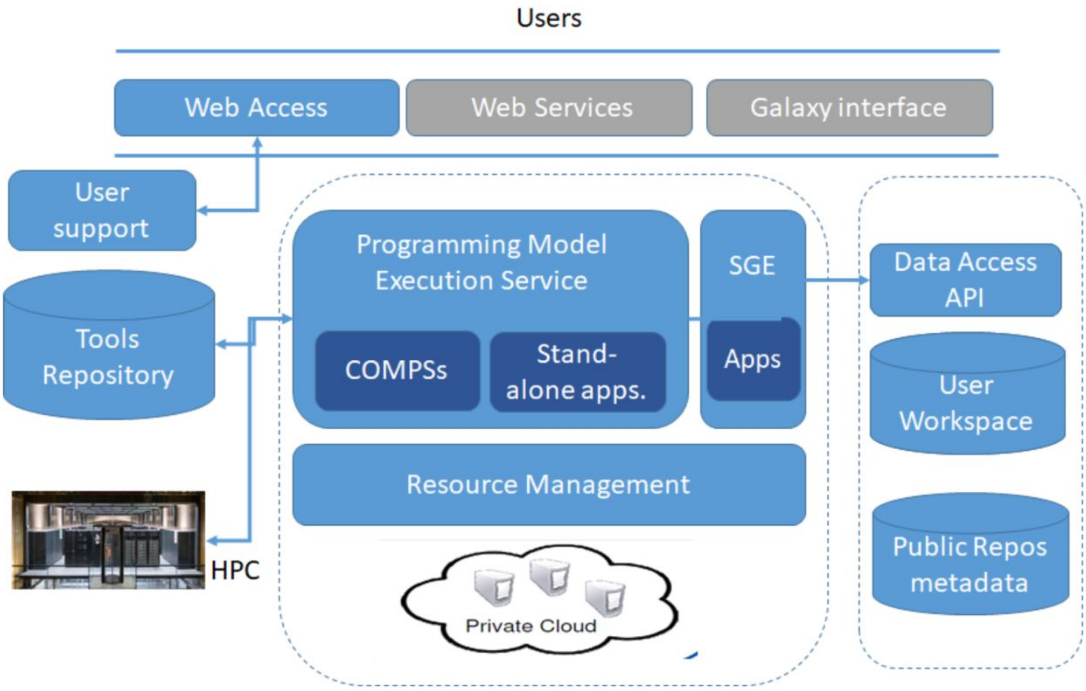


**Figure S1.** Scheme of MuGVRE computational infrastructure.
Greyed modules will be available by the end of 2019

#### *MuGVRE Main components*

*Cloud deployments:* MuGVRE infrastructure has been designed as a fully virtualized environment. This layout allows to deploy instances of the VRE Backend in new cloud infrastructures with minimal overhead. Besides, the deployment of tools as virtual machines allows to configure an elastic infrastructure, to cover peaks of demand, or to configure complex workflow schemes. MuGVRE has been deployed at the StarLife cloud infrastructures, at the Barcelona Supercomputing Center, using OpenNebula (https://opennebula.org/) and the KVM hypervisor (https://www.linux-kvm.org). However, the generation of Virtual Machines has been adapted to make them compatible with the deployment in both openNebula and openStack (https://www.openstack.org/) cloud managers, allowing their use in a wider set of cloud platforms, including Elixir Compute Platform (http://elixir-europe.eu) and EGI (https://egi.eu) providers.

*Process management:* MuGVRE uses two complementary options for process management:

1) Sun Grid Engine (SGE, https://sourceforge.net/projects/gridscheduler/), in combination with OneFlow (https://docs.opennebula.org/5.4/advanced_components/application_flow_and_auto-scaling/appflow_use_cli.html), a component of the OpenNebula framework that allows

managing Multi-VM application and auto-scaling. SGE is used to manage applications where no complex workflows are necessary, requiring only to deploy additional workers on peaks of demand. Each execution is sent to a tool specific SGE queue that can be populated with multiple instances of the VM where the tool is packaged. OneFlow controls the number of running instances, deploying then dynamically according to a set of configurable system metrics.

2) The COMPS Superscalar (COMPSs) (1) programming model, combined with programming Model Enactment Service (PMES) (2). COMPSs programming model and runtime are designed to simplify the development and execution of distributed applications. COMPSs applications are programmed in a completely sequential manner but contain code annotations to define data dependencies and execution requirements at the task level. Using these annotations, COMPSs runtime can automatically detect and exploit the inherent parallelism of the application, and execute it on distributed platforms, such as Grids, Clouds, and clusters. In the case of elastic infrastructures such as Clouds, the runtime can dynamically create and destroy workers to tailor the computational capacity to the application workload. COMPSs runtime implements a master-worker architecture (see Suppl. Figure S2). Master and workers can run on different virtual machines (VM) or physical nodes depending on the characteristics of the underlying infrastructure. In the context of the MuGVRE, COMPSs Python binding (also known as PyCOMPSs) is used due to familiarity with this programming language in the bioinformatics field. The Programming Model Enactment Service (PMES) controls the execution of jobs in an underlying Cloud platform through an Open Cloud Computing Interface (OCCI, http://occi-wg.org/) Server. Although PMES supports single command jobs, it is used mainly to control COMPSs application using one or more VMs, allowing the execution of large parallel workflows. The OCCI Server(s) thus abstract the PMES and COMPSs runtime from the underlying infrastructure and allows the execution of applications using any OCCI compliant Cloud middleware, in particular, OpenNebula or OpenStack.

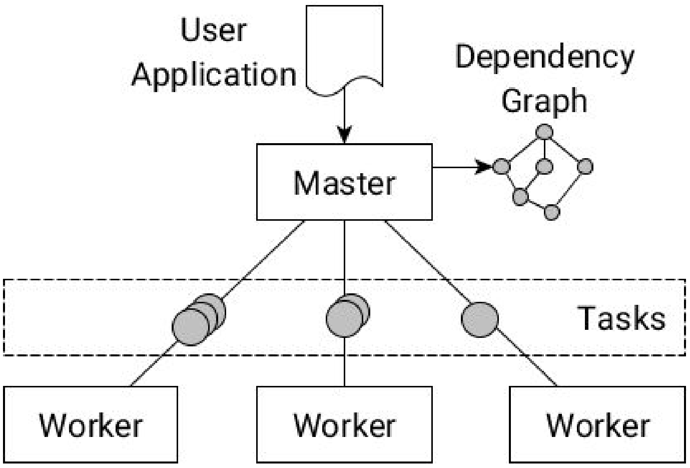


**Figure S2.** COMPSs master-worker architecture.

*Database managers:* MuGVRE data is divided in two types of repositories. A MongoDB database (https://www.mongodb.com) is used to hold operational data (user management, job execution, sample data collection, help) and metadata corresponding to user files, and to installed tools. This set of metadata is used to correctly combine data with the appropriate tools and visualizers (see Section 2 at this Suppl. Material for a summary of this process, and the data models used). The MongoDB server where MuG data is hosted, contains also reference data as a full copy of Protein Data Bank (3) , and Uniprot (4), and the trajectory database BiGNASim (5); this eases access to this information from the workspace. Data itself is stored in a standard filesystem in its original format. The filesystem layout is organized per user (including anonymous users) so that the privacy of data is maintained in all cases.

*Integration of remote repositories:* MuG aims to ease the access of users to relevant public data repositories, where studies related to 3D/4D genomics are being maintained. In the present version of MuGVRE, metadata from selected studies of BigNASim(5) and ArrayExpress (6) have been stored in the MongoDB metadata repository. Metadata allows users to browse and search for specific studies using MuGVRE interface and download data into the personal workspace for further analysis.

### PROCEDURE FOR THE MANAGEMENT OF TOOLS IN MuGVRE

The modular and portable design of the MuGVRE computational platform has led to the complete virtualization of the tools and pipelines. Tools live encapsulated in virtual machines, and the VRE core acts as a framework that delivers to them the input files and their metadata, sets up the deployment procedure of the VMs, monitors tool execution, and eventually gathers the output files and their metadata once the execution has finished. To perform all these procedures, a protocol defining how MuGVRE core communicates with the virtualized tools has been established. The following sections describes i) how tool developer is to prepare a virtualized VRE tool, and ii) how VRE manages a tool execution.

### Integration of a new tool

The metadata for the tools integrated in MuGVRE is stored in a MongoDB. This metadata includes:

(i) *descriptive data,* used to illustrate the tool inside the VRE and automatically build its help section,

(ii) *deployment details,* like the required computational resources, the process manager in use (SGE or PMES), the remote path where the application is the be called, or the cloud infrastructure where the virtualized application is installed.

(iii) *a definition of the input and output files* in terms of data type and format (table S3), as well as other arguments the application may take.

With this information VRE learns how to adequately manage the tool:

- Suggest the tool given a set of input files in the user workspace
- Create a web form so that the user can specify the arguments before execution
- Invoke the application via any of the process managers (See next section)
- Monitor the tool progress during an execution and detecting when an error occurred
- Register the tool results in the Data Management protocol (DMP) so output files can be found in the user's workspace.
- Recognize the ownership of the tool, so that tool developers have the adequate administrative permissions over their tools

To incorporate a new tool into the system, the developer needs to prepare, along with the tool VM itself, a *tool definition* file (JSON format) containing such information. The JSON schema and an example of *tool definition* file are available in Table S1.

**Table S1.** Schema and example of Tool definition configuration file, required for registering a new tool

| |
|---|
| 1. Tool definition (JSON – schema)<br>https://github.com/Multiscale-Genomics/VRE_tool_jsons/blob/dev/tool_specification/tool_schema.json |
| 2. Tool definition (JSON – example) |

https://github.com/Multiscale-Genomics/VRE_tool_jsons/blob/dev/tool_specification/examples/pydockdna.json

For users registered as tool developers, VRE includes an online section under the "admin" menu for guiding and helping on the *tool definition* file generation. Is is based on the JSON schema validation, and includes the generation of test files that allow to emulate a VRE call in the development environment where the user is working. After tests are passed, a ticketing-based dialog is open between the tool developer and the VRE administrators before the tool VRE is published in the infrastructure. An step-by-step documentation is available here: https://www.multiscalegenomics.eu/MuGVRE/instructions/

### Tool execution lifecycle

Once the tool is properly defined, it is ready to be launched by the MuGVRE execution engine. Figure S3 covers the complete life cycle of a tool execution and summarizes the data flow carried out in each step.

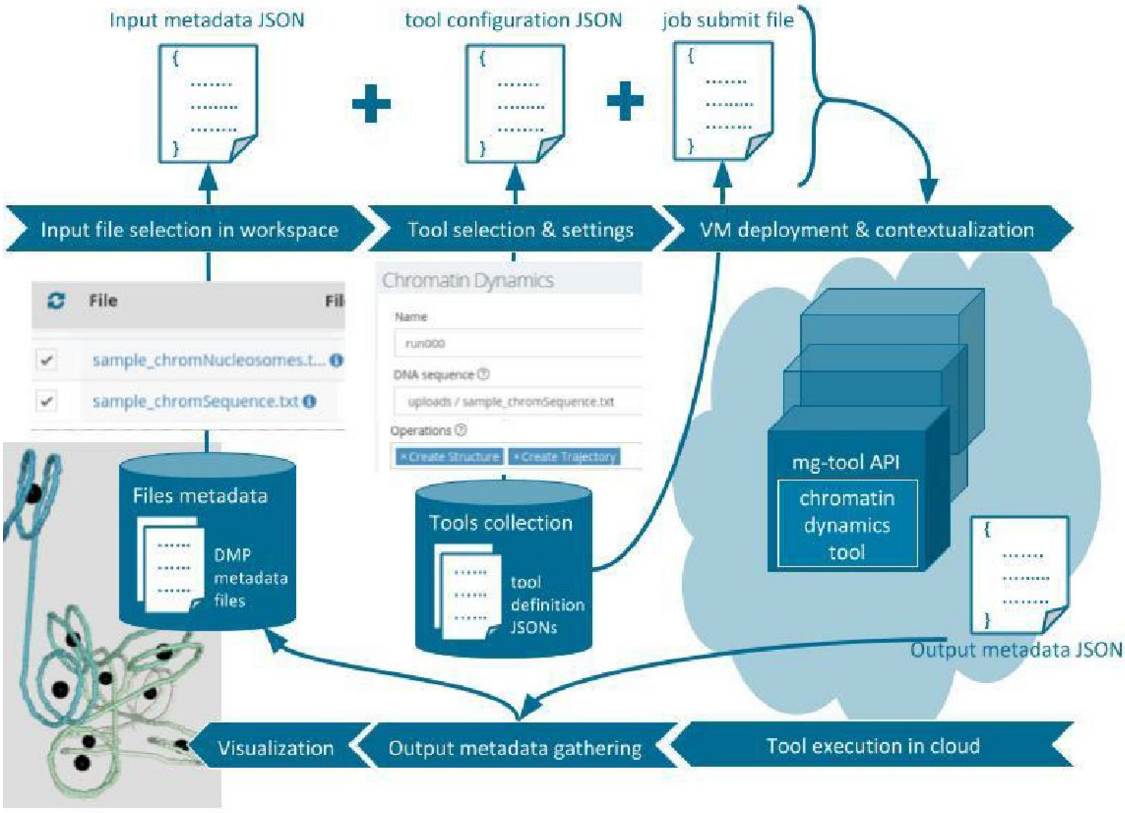

**Figure S3**. Life cycle of a tool execution in MuGVRE, and how the information is transferred from the VRE user to the virtualized tools, and back to the VRE user.

The end user defines the input files and arguments a new execution on a web form created mostly automatically based on the Tool definition. Such information is transferred to the tool via two JSON files called *input metadata* and *tool configuration* (examples in Table S2). The first contains the metadata corresponding to the input files, including data types and formats, as well as, the disk path as it is to be seen by the virtualized environment. The second file contains the parameter values for the particular run. When the user clicks the "Compute" button, and according to the *tool definition* JSON, one of the two process managers supported by the VRE (PMES, SGE-oneflow) is triggered. In short, if SGE-oneflow is the election, a *submit* bash file with the application command line will be submitted to a queue hosted a the tool virtual machine (example in Table S2). If a pick on demand occurs, (the tool VM load is persistently high during certain time and exist a number of pending execution), onFlow detects so and responds deploying extra instances of that virtual machine, offering extra computation resources that will be undeployed after the pick demand. If PMES is the selected, a REST call to the *create activity* endpoint (2) will be performed, and the tool VM will be deployed, contextualized, to eventually execute the application command line.

**Table S2:** Examples for the configuration files exchanged between VRE and tool VMs during the tool life cycle execution

| |
|---|
| 3. Input metadata (JSON – example)<br>https://github.com/Multiscale-Genomics/VRE_tool_jsons/blob/dev/tool_execution/sample_project/myPydockProject/.input_metadata.json |
| 4. Configuration tool (JSON – example)<br>https://github.com/Multiscale-Genomics/VRE_tool_jsons/blob/dev/tool_execution/sample_project/myPydockProject/.config.json |
| 5. Submit file - examples<br>https://github.com/Multiscale-Genomics/VRE_tool_jsons/blob/dev/tool_execution/sample_project/myPydockProject/.submit |
| 6. Output metadata (JSON – example)<br>https://github.com/Multiscale-Genomics/VRE_tool_jsons/blob/dev/tool_execution/sample_project/myPydockProject_out/.results.json |

If the application is to be run under pyCOMPSs, the process manager will invoke the application executable using pyCOMPSs libraries. The application itself in general inherits from mg-tool API (https://github.com/Multiscale-Genomics/mg-tool-api), which implements a homogenous layer on top of the application code that both, transparently deals with communication to and from the MuGVRE core, and absorbs possible application heterogeneities.

Once the execution is finished, the last step performed by the MuGVRE engine is to gather the tool output files. Metadata to qualify result files is taken from the *tool definition* JSON. When some (or all) of the details of the output files cannot be in advance (e.g. when then depend on the inputs), each execution of the tool should create an additional JSON file, called *output metadata,* (see Table S2), which contains the details of the specifics outputs.

### Data type and formats registered at MuGVRE

To be able to automatically decide on the compatibility of tools and data, a comprehensive analysis of the tools available and their requirements has led to the building of an ontology of data types and formats (see Table S3). Data types and formats are always managed as a single unit defining a series of possible combinations. The procedure to select tools and visualizers considers the number and type of parameters accepted by the tool and generates a list of allowed combinations for each tool that are then matched against user files. Metadata for user files is provided by the user (when it cannot be guessed from the file format) for uploaded data or taken from the *tool definitions* JSON file.

**Table S3. Data types and formats**

| Data type | Associated file type (format) |
|---|---|
| 3D structure | PDB |
| ATAC-Seq | FASTQ, BAM, BED, WIG |
| BWA index files | AMB, ANN, BWT, PAC, SA, TAR |
| Bowtie2 index files | BT2, TXT, TAR |
| ChIP-Seq | BED, FASTQ, BAM, TSV |
| Chromatin 3D structure | PDB |
| Chromatin TADs | BED, TXT |
| Chromatin compartments data | TXT |
| Chromatin trajectory | DCD |
| DNA methylation | FASTQ, WIG, TSV, BW |
| DNA sequence | FASTA, TXT |
| Docking ranking score | CSV, TXT, TSV |

| | |
|---|---|
| Ensemble of chromatin 3D structures | JSON |
| FISH data | LIF, TIFF, PNG |
| Genomic sequence | FASTA |
| HiC Biases | PICKLE |
| HiC TADs scaling factor | WIG |
| HiC aligned reads | BAM |
| HiC contact matrix | TXT, HDF5 |
| HiC contact peaks | TSV |
| HiC contacts coverage | WIG, BW, TXT |
| HiC differential contacts | TSV |
| HiC directionality index | TXT |
| HiC sequencing reads | FASTQ |
| Kallisto index file | IDX |
| MD restart file | RST, CPT |
| MNase-Seq | FASTQ, BAM, BED |
| Nucleic acid 3D structure | PDB |
| Nucleic acid MD CG trajectory | MDCRD |
| Nucleic acid MD atomistic trajectory coordinates | XTC, NETCDF, MDCRD |
| Nucleic acid MD atomistic trajectory topology | TOP, TPR, PARMTOP, PDB |
| Nucleic acid topology | TOP, TPR, PARMTOP, PDB |
| Nucleic acid trajectory | DCD, MDCRD |
| Nucleic acid trajectory coordinates | XTC, NETCDF, MDCRD |
| Nucleosome TSS | BW, GFF3, BED, WIG |
| Nucleosome dynamics | BW, GFF3, BED, WIG, RDATA |
| Nucleosome free regions | BW, GFF3, BED, WIG |
| Nucleosome phasing | BW, GFF3, BED, WIG |
| Nucleosome positioning | BW, GFF3, BED, WIG, TXT |
| Nucleosome stiffness | BW, GFF3, BED, WIG |
| Protein 3D structure | PDB |
| Protein sequence | FASTA |
| Protein-DNA complex structure | PDB |
| Protein-DNA specificity | TSV |

| | |
|---|---|
| Protein-Protein complex structure | PDB |
| RNA sequence | FASTA |
| RNA-Seq | FASTQ, TSV, HDF5, JSON, BAM, BW, GFF, GFF3 |
| Sequence Annotation | BED, BB, BEDGRAPH, WIG, BW, GFF, GFF3, GTF, VCF, TBI |
| Sequence mapping index | GEM |
| Tool Intermediate file | TAR |
| Tool configuration file | JSON, TXT, TSV |
| Tool summary file | TAR |
| Whole Genome Bisulfite Sequencing | FASTQ, BAM, BAI, WIG, TSV, TXT |
| cDNA sequence | FASTA |

### Analysis and simulation tools available

Tools incorporated within MuGVRE cover many levels of the analysis of 3D/4D genomics. The portfolio of offered tools is expected to expand, as the infrastructure provide a straightforward procedure to implement new tools, in particular requiring little or no modification of the original software (see above). The following table shows all the tools currently available within MuGVRE.

**Table S4.** Analysis and Simulation tools

The present table summarizes the tools integrated or in the process of being integrated in MuGVRE together with some of their implementation details.

| Tool name | Category | Description | Implementation |
|---|---|---|---|
| Bowtie2 | -DNA | Align FASTQ data using Bowtie2 | Manager: PMES<br>Skeleton: mg-tool API<br>Job type: PyCOMPSs |
| BWA MEM | -DNA | Align FASTQ data using BWA mem | Manager: PMES<br>Skeleton: mg-tool API<br>Job type: PyCOMPSs |
| Chromatin Dynamics | -Chromatin<br>-DNA | Chromatin Dynamics provides a user-friendly way to create individual 'beads-on-a-string' like | Manager: SGE-oneflow<br>Skeleton: custom wrapper<br>Job type: single |

| | | | |
|---|---|---|---|
| | | representations of a chromatin fiber. | |
| MACS2 | -DNA | MACS identifies statistically significantly enriched genomic regions in ChIP- and DNase-seq data | Manager: PMES<br>Skeleton: mg-tool API<br>Job type: PyCOMPSs |
| MC-DNA | -DNA | MC-DNA is a tool to rapidly create static and dynamic B-DNA conformations of a sequence of interest. With the use of a Monte Carlo algorithm this tool runs up to 50x faster than conventional Molecular Dynamics providing similar accuracy.<br>MC-DNA provides a three-dimensional all-atom representation of the DNA structure with the underlying sequence of interest. | Manager: SGE-oneflow<br>Skeleton: custom wrapper<br>Job type: single |
| MDWeb<br>(MD Energy refinement) (8) | -DNA<br>-Protein<br>-RNA | MDWeb is based on well-known simulation programs like Amber, NAMD and Gromacs, and a series of preparation and analysis tools, joined together in a common interface. | Manager: SGE-oneflow<br>Skeleton: custom wrapper<br>Job type: single |
| NAFlex (9) | -DNA<br>-RNA | NAFlex provides a friendly environment to analyse your own generated molecular dynamics trajectories of nucleic acid structures. | Manager: SGE-oneflow<br>Skeleton: custom wrapper<br>Job type: single |
| Nucleosome Dynamics | -DNA | Nucleosome positioning plays a major role in transcriptional regulation and most DNA-related processes. The nucleosome dynamics server offers different tools to analyze nucleosome positioning from MNase-seq experimental data and perform comparative | Manager: SGE-oneflow<br>Skeleton: custom wrapper<br>Job type: single |

| | | | |
|---|---|---|---|
| | | experiments to account for the transient and dynamic nature of nucleosome positioning under different cellular states. | |
| 3D Consensus | -DNA<br>Interaction s<br>-Protein | Analyze a protein-DNA complex 3D structure to identify interactions and study their impact on specific binding by integrating experimental data on the protein's DNA specificity. 3DConsensus allows the interpretation of experimental data on DNA-binding specificity of a protein through the analysis of a 3D structure of the complex. | Manager: SGE-oneflow<br>Skeleton: mg-tool API<br>Job type: single |
| Process Genome (https://github.com/Multiscale-Genomics/mg-process-fastq) | -DNA | Pipeline for generating index files for a genomic sequence. Once the index files have been generated for a given assembly then they can be used by different pipelines/tools as they are required. Based on the FASTA file of a genomic sequence index files are generated for the following indexers: Bowtie2, BWA, GEM | Manager: PMES<br>Skeleton: mg-tool API<br>Job type: PyCOMPSs |
| pyDock (10) | -Interaction s<br>-Protein | pyDock is a tool for the structural prediction of protein-protein interactions. | Manager: SGE-oneflow<br>Skeleton: custom wrapper<br>Job Type: single |
| pyDockDNA (https://github.com/Multiscale-Genomics/pydockdna_tool) | -DNA<br>-Interaction s<br>-Protein | pyDockDNA is a tool for the structural prediction of protein-DNA interactions. | Manager: SGE-oneflow<br>Skeleton: custom wrapper<br>Job type: single |
| TADBit (11) | -Chromatin<br>-DNA | TADbit is a complete Python library to deal with all steps to analyze, model and explore 3C-based data | Manager: SGE-oneflow<br>Skeleton: custom wrapper<br>Job type: single |
| TADBit map filter and parse | | Maps and filters Hi-C read FASTQ files obtains a | |

| | | | |
|---|---|---|---|
| | | pseudo BAM with the aligned reads. | |
| TadBit Normalize | | Normalize the aligned Hi-C reads. | |
| TadBit Segment | | finds Topologically Associating Domains (TAD)s and segments | |
| TadBit Binning | | bins interaction matrices. | |
| TadBit Modeling | | builds an ensemble of 3D models from the interaction matrices, able to be explored and visualized in TADkit. | |
| Process ChiP-seq (https://github.com/Multiscale-Genomics/mg-process-fastq) | -DNA | Pipeline for processing ChIP-seq sequence reads to identify regions of DNA-protein interactions. Sequences are aligned to the genomic sequence using BWA, BioBamBam2 is used to filter out experimental artifacts and MACS2 is used for the analysis of the alignments to identify regions of DNA-protein interaction. | Manager: PMES<br>Skeleton: mg-tool API<br>Job type: PyCOMPSs |
| Process RNA-seq (https://github.com/Multiscale-Genomics/mg-process-fastq) | -RNA | Align RNA-seq data pipeline. Gene expression calling with Kallisto. | Manager: PMES<br>Skeleton: mg-tool API<br>Job type: PyCOMPSs |
| Process WGBS (https://github.com/Multiscale-Genomics/mg-process-fastq) | -DNA | Align WGBS (Whole-Genome Bisulfite Sequencing) data. Uses BS Seeker2 and Bowtie2 | Manager: PMES<br>Skeleton: mg-tool API<br>Job type: PyCOMPSs |

**Table S5.** Data visualizers available at MuG VRE

| Visualizer | Description | Supported Data |
|---|---|---|
| NGL Viewer (12) | NGL Viewer is a web application for 3D molecular structure visualization | 3D Structures and MD trajectories (PDB, DCD) |
| JBrowse (13) | JBrowse is a fast, embeddable genome browser built completely with JavaScript and HTML5. | Genome sequence annotation related formats (BAM, BW, GFF, GFF3) |

| | | |
|---|---|---|
| TADKit<br>http://3DGenomes.org/tadkit | TADkit creates interactive 3D representations of chromatin conformations modeled from 3C-based interaction matrices, overlaid with 1D and 2D tracks of genomic data. | HiC analysis data processed with TADBit<br>(JSON, TXT) |

### EXAMPLE OF USE. NUCLEOSOME DYNAMICS

This overview shows the use of MuGVRE interface to perform a Nucleosome Dynamics analysis on MNase-seq data of yeast chromosome II at two phases of the cell cycle (G2 and M). Data is taken from the study by Deniz *et al*. (14). Experiments correspond to BY4741 yeast cells, synchronized at late G1 using alpha-factor pheromone. Samples were collected at 45' (G2) and 60' (M), digested with MNase, and sequenced with Illumina HiSeq 2000. Data is available at the MuGVRE workspace as Sample Data. as BAM files (cellcycleG2_chrii.bam and cellcycleM_chrii.bam), aligned against Yeast R64-1-1 genome assembly. Figure S4 show the validation form where the used should indicate the data type, and Taxon Id. Initial values can be guessed by the system, but the data type "MNase DNA seq" should be included by the user to distinguish from other types of sequenced data. Validation also make a quality control of the uploaded files. Figure S5 shows the upload folder after data upload and validation. Toolkits corresponding to BAM files are shown.

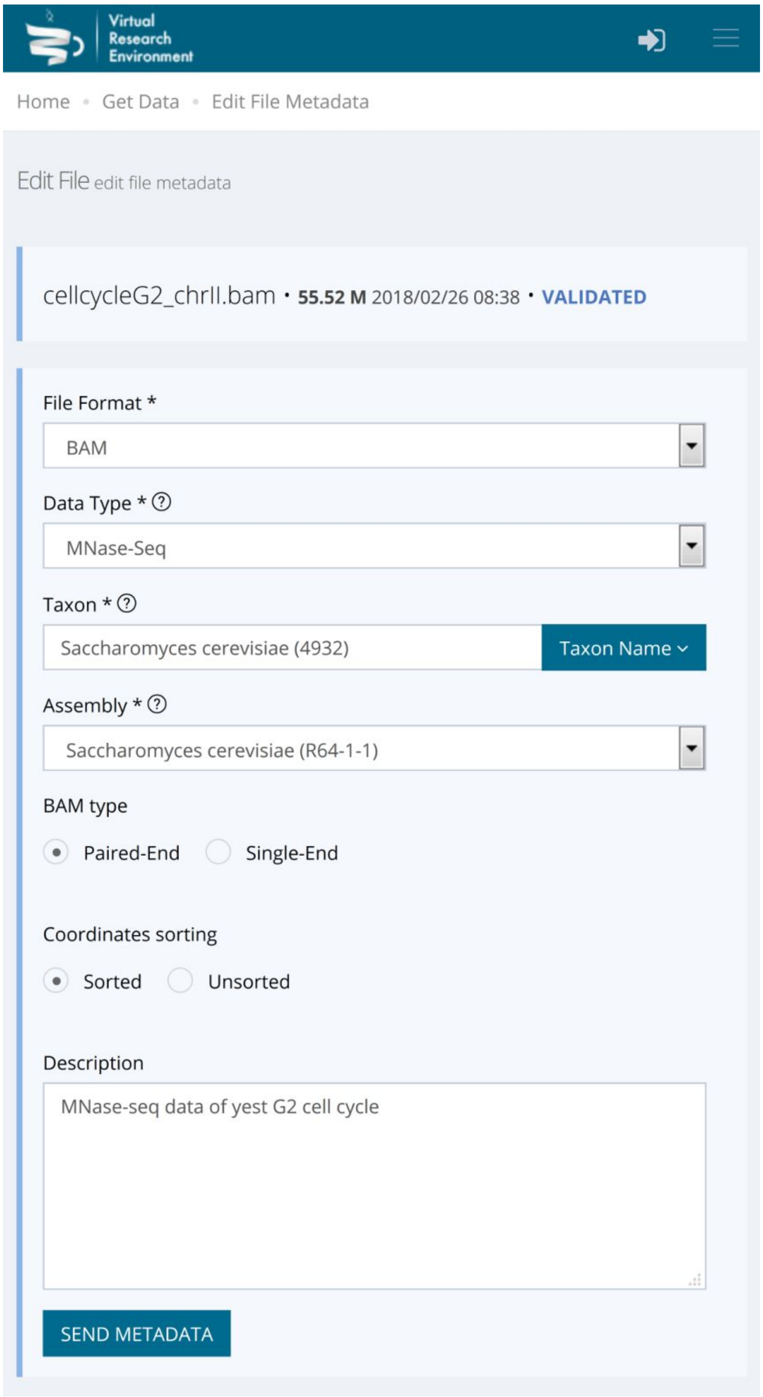


**Figure S4**. Validation Form required to add the necessary metadata to uploaded files.

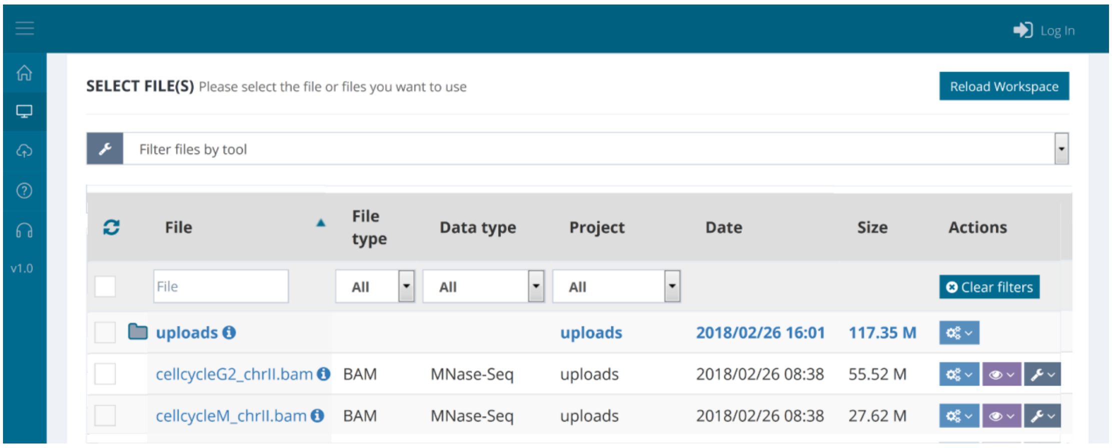


**Figure S5:** Workspace with toolkits

At this point input file can be examined. Figure S6 shows a JBrowse view of the sequence coverage of input data

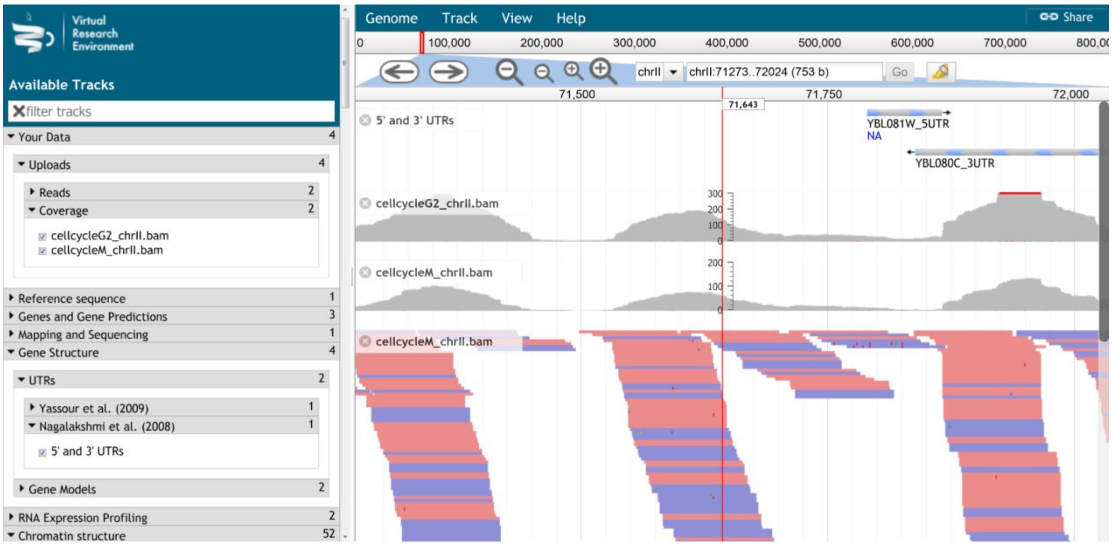


**Figure S6.** JBrowse view of coverage of input BAM files

Nucleosome Dynamics can perform a series of operation on this data: 1) Nucleosome Calling, 2) Nucleosome Free Regions, 3) Nucleosome dynamics, showing differences between the two experimental cases, 4) Nucleosome Phasing, 5) Classification of TSS, 6) Evaluation of stiffness in nucleosome attachment. When selecting the tool, the user is given the opportunity of selecting the operations to perform and qualify the input files according to the tools requirements (Figure S7). After this, the calculation can be submitted.

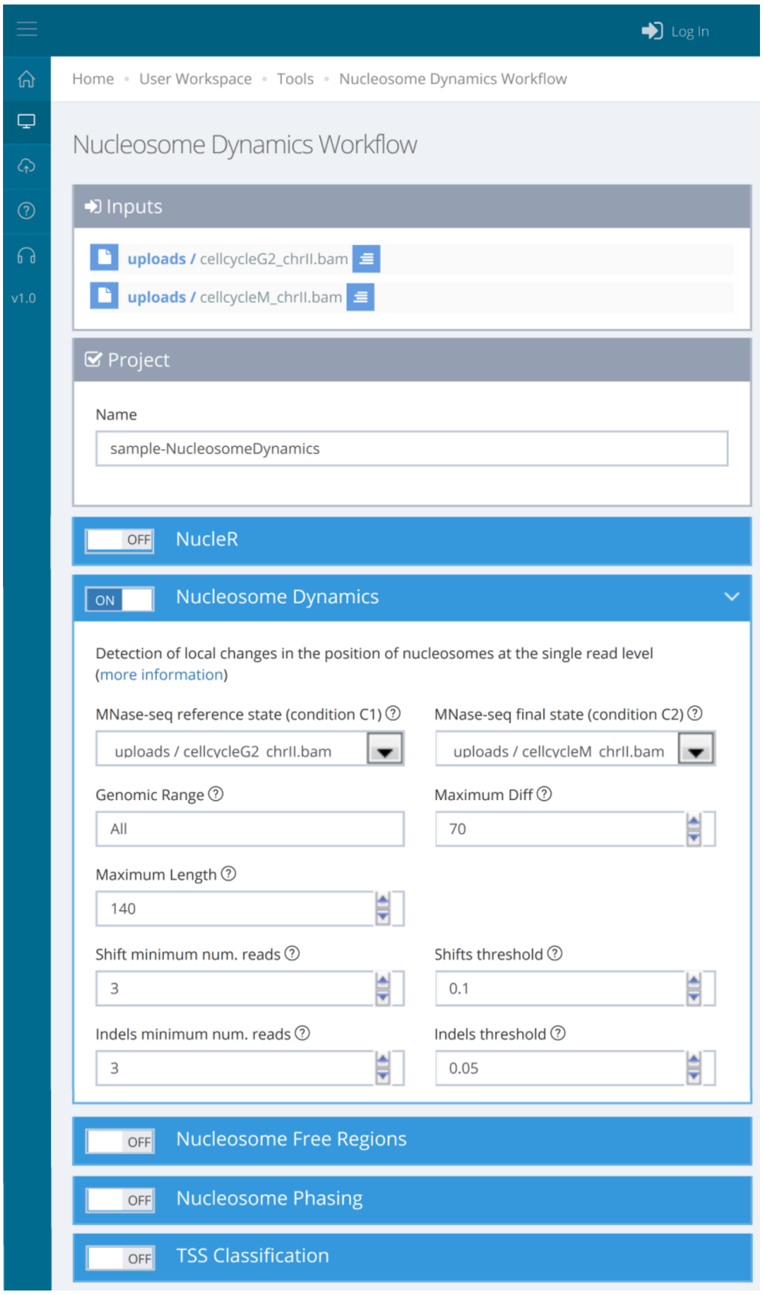


**Figure S7**: Parameter input and submit

During tool execution, VRE keeps track of the job progress indicating the queue or deployment status (PENDING, BOOT, EPILOG, RUNNING, FINISHING, ERROR) at the user's workspace. While RUNNING, an updated tool log is displayed either, in the raw format, as the application in execution returns, or in a digested view, where the raw log is formatted for clearance and better understanding.

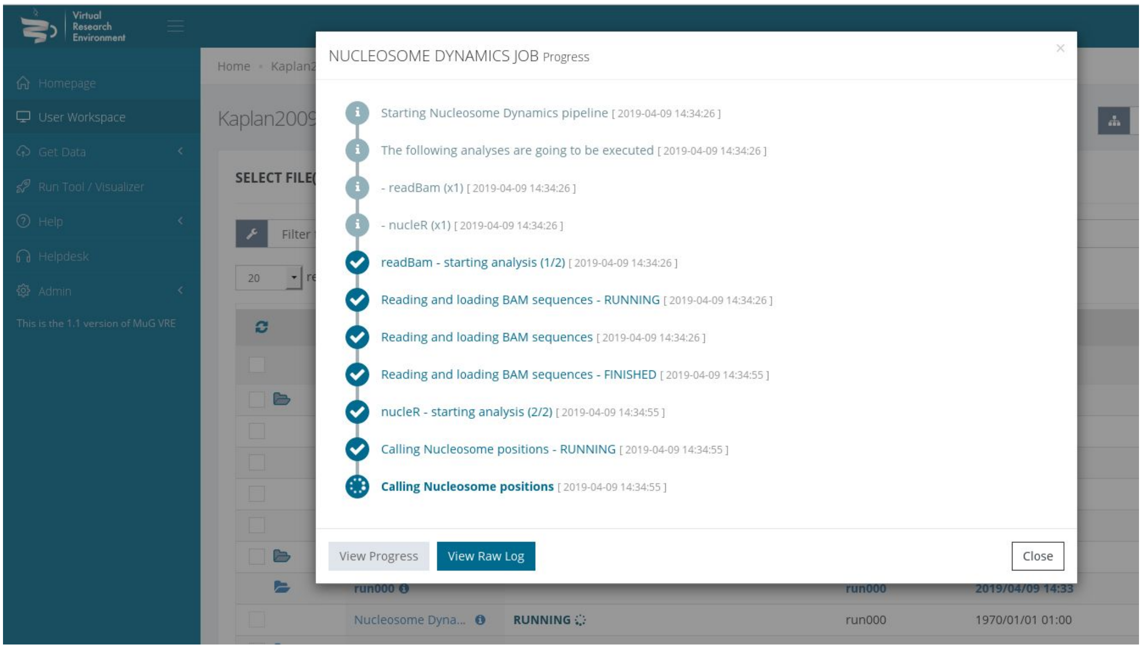


**Figure S8**: Job progress display. Digested view of the application log

After completion, results are incorporated to the workspace and can be analyzed (Figure S8).

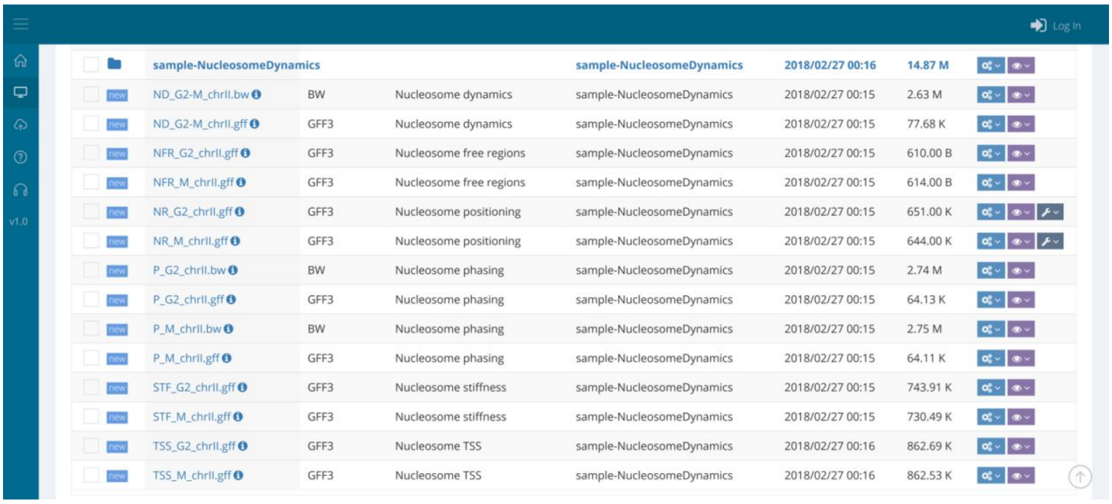


**Figure S9**: Project folder including output files

A specific results page is prepared showing aggregated results (Figure S10). Individual output data files can be analyzed. See Figure S11 for a screenshot of JBrowse showing the results of Nucleosome Dynamics analysis

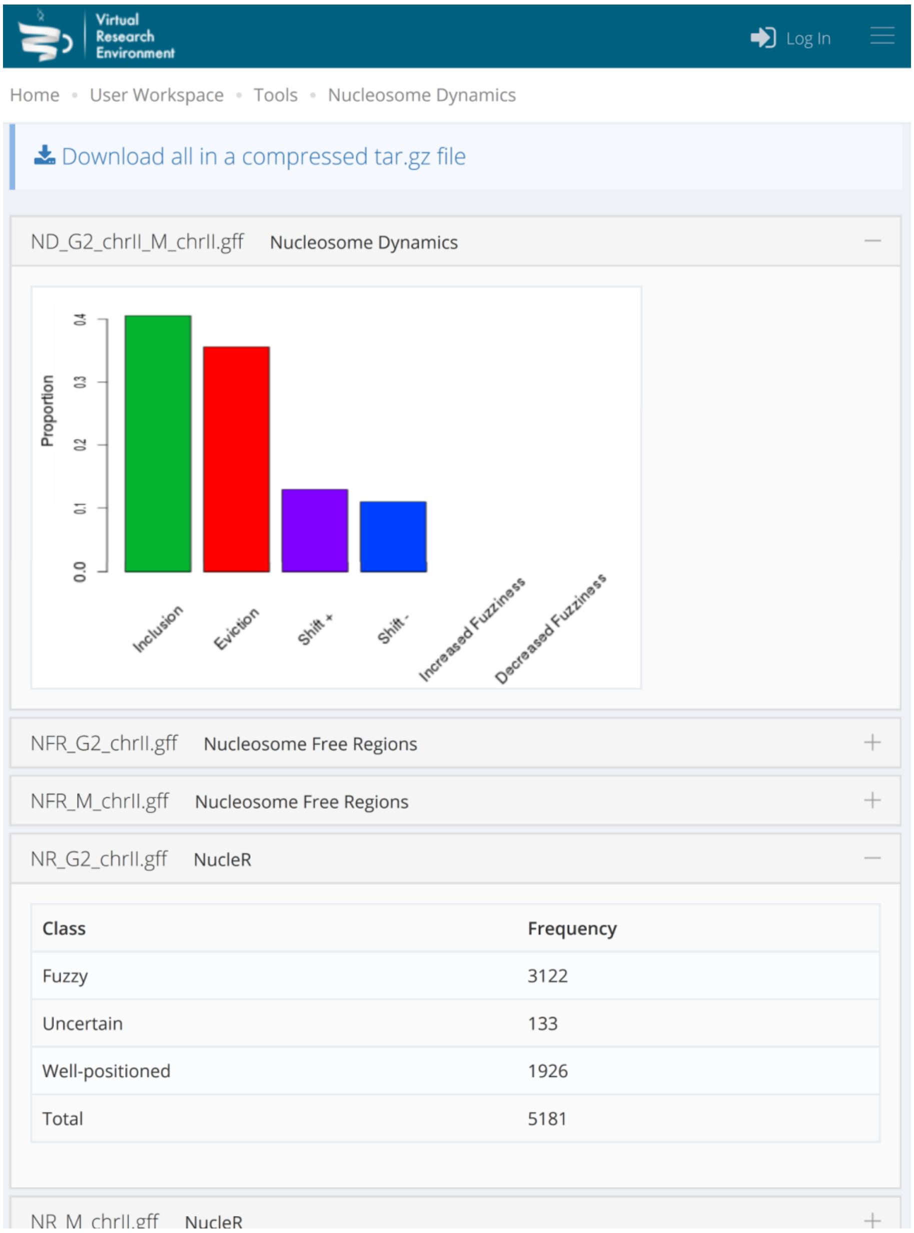


**Figure S11** Summary results page

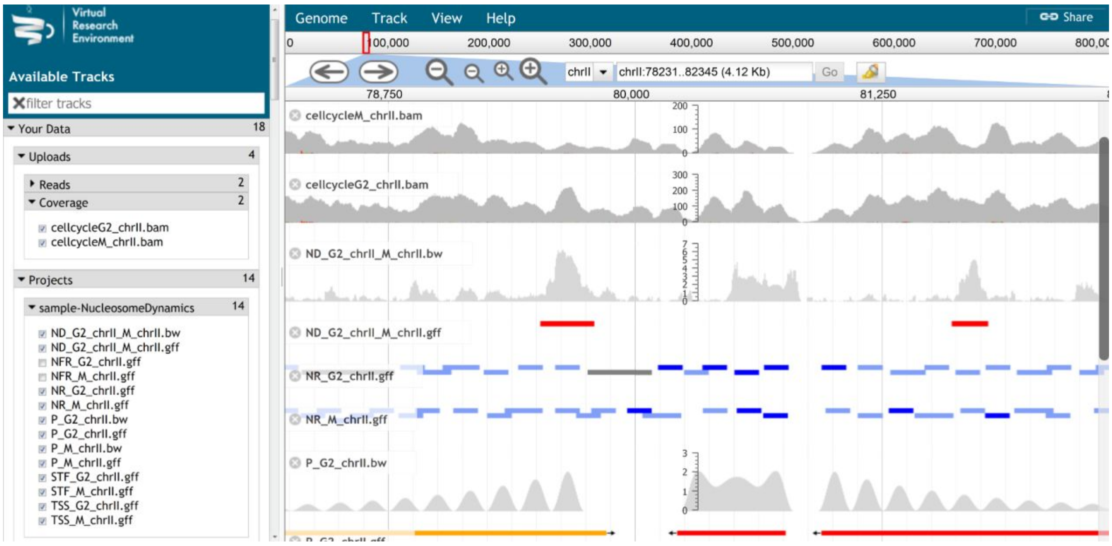


**Figure S11.** JBrowse view of Nucleosome analysis results

Some of the output files can be further analyzed. For instance, nucleosome position called can be transferred and simulated using the Chromatin Dynamics tools to gather the dynamics of the DNA fiber including the effect of nucleosomes.
